## Supplemental Table 1 for "Large vesicle extrusions from *C. elegans* neurons are consumed and stimulated by glial-like phagocytosis activity of the neighboring cell"

| Strain name | Genotype | Reference |
| --- | --- | --- |
| ZB4690 | *sid-1(qt9); bzIs101[pmec-4::mCherry]; bzSi13[phyp-7::sid-1]* | [7] |
| ZB4757 | *bzIs166[pmec-4::mCherry]* | [7] |
| RT4491 | *uIs31[mec-17p::GFP]* | [66] |
| RT3870 | *bzIs166[pmec-4::mCherry]; pwSi144[phyp-7::mNeonGreen::lgg-1]* | This study |
| RT3835 | *bzIs166[pmec-4::mCherry]; pwSi140[phyp-7::mNeonGreen::rab-7]* | This study |
| RT4090 | *bzIs166[pmec-4::mCherry]; pwSi226[phyp-7::mNeonGreen::rab-5]* | This study |
| RT3793 | *bzIs166[pmec-4::mCherry]; pwSi107[phyp-7::mNeonGreen::ced-10]* | This study |
| RT3818 | *bzIs166[pmec-4::mCherry]; pwSi130[phyp-7::mNeonGreen::2XFYVE]* | This study |
| RT3824 | *bzIs166[pmec-4::mCherry]; pwSi132[phyp-7::mNeonGreen::rme-1]* | This study |
| RT3834 | *bzIs166[pmec-4::mCherry]; cnt-1(tm2313)* | This study |
| RT3855 | *bzIs166[pmec-4::mCherry]; rab-35(b1034)* | This study |
| RT3930 | *bzIs166[pmec-4::mCherry]; pwSi178[phyp-7::mNeonGreen::rab-35]; rab-35(b1034)* | This study |
| RT3929 | *bzIs166[pmec-4::mCherry]; pwSi177[pmec-7::mNeonGreen::rab-35]; rab-35(b1034)* | This study |
| RT3934 | *bzIs166[pmec-4::mCherry]; pwSi181[pmec-7::cnt-1::mNeonGreen]; cnt-1(tm2313)* | This study |
| RT3937 | *bzIs166[pmec-4::mCherry]; pwSi184[phyp-7::cnt-1::mNeonGreen]; cnt-1(tm2313)* | This study |
| RT3995 | *bzIs166[pmec-4::mCherry]; pwSi202[phyp-7::aman-2(1-82)::mNeonGreen]* | This study |
| RT3998 | *bzIs166[pmec-4::mCherry]; pwSi205[phyp-7::lmp-1::mNeonGreen]* | This study |
| RT4107 | *bzIs166[pmec-4::mCherry]; pwEx63[phyp-7::PH plcΔ::mNeonGreen]* | This study |
| RT4136 | *bzIs166[pmec-4::mCherry]; arf-6(tm1447)* | This study |
| ZB5211 | *bzIs166[pmec-4::mCherry]; arf-6(ns388)* | This study |
| RT4643 | *uIs31[pmec-17::GFP]; arf-6(tm1447)* | This study |
| RT4140 | *bzIs166[pmec-4::mCherry]; pwSi205[phyp-7::lmp-1::mNeonGreen]; cup-5(ar465)* | This study |
| RT4142 | *bzIs166[pmec-4::mCherry]; zdIs5[pmec-7::GFP]; ced-1(e1735)* | This study |
| RT4614 | *bzIs166[pmec-4::mCherry]; zdIs5[pmec-7::GFP];  pwSi504[Phyp7::CED-1::GFP]; ced-1(e1735)* | This study |
| ZB5275 | *bzIs166[pmec-4::mCherry]; ced-1(e1735)* | This study |
| ZB4086 | *bzIs166[pmec-4::mCherry]; zdIs5[pmec-7::GFP]* | [7] |
| RT4180 | *bzIs166[pmec-4::mCherry]; pwSi184[phyp-7::cnt-1::mNeonGreen]; rab-35(b1034)* | This study |
| RT4181 | *bzIs166[pmec-4::mCherry]; pwSi178[phyp-7::mNeonGreen::rab-35]; cnt-1(tm2313)* | This study |
| RT4183 | *bzIs166[pmec-4::mCherry]; arf-6(tm1447); cnt-1(tm2313)* | This study |
| RT4184 | *bzIs166[pmec-4::mCherry]; arf-6(tm1447); rab-35(b1034)* | This study |
| RT4185 | *bzIs166[pmec-4::mCherry]; pwSi229[phyp-7::arf-6::mNeonGreen]; arf-6(tm1447)* | This study |
| RT4178 | *bzIs166[pmec-4::mCherry]; pwSi229[phyp-7::arf-6::mNeonGreen]; cnt-1(tm2313)* | This study |
| RT4097 | *bzIs166[pmec-4::mCherry]; pwSi229[phyp-7::arf-6::mNeonGreen]* | This study |
| RT4231 | *bzIs166[pmec-4::mCherry]; arf-6(tm1447); daf-2(e1370)* | This study |
| ZB4857 | *bzIs166[pmec-4::mCherry]; daf-2(e1370)* | [7] |
| RT4273 | *bzIs166[pmec-4::mCherry]; pwSi369[phyp-7::UtrCH::mNeonGreen]* | This study |
| RT4414 | *bzIs166[pmec-4::mCherry]; pwSi406[phyp-7::ced-1Δ::GFP]* | This study |
| RT4404 | *bzIs166[pmec-4::mCherry]; pwSi369[phyp-7::UtrCH::mNeonGreen]; sand-1(or552)* | This study |
| RT4416 | *bzIs166[pmec-4::mCherry]; arl-8(wy271)* | This study |
| RT4489 | *bzIs166[pmec-4::mCherry]; ced-10(n3246)* | This study |
| RT4571 | *bzIs166[pmec-4::mCherry]; ttr-52(tm2078)* | This study |
| RT4570 | *bzIs166[pmec-4::mCherry]; anoh-1(tm4762)* | This study |
